# Functional classes of SNPs related to psychiatric disorders and behavioral traits contrast with those related to neurological disorders

**DOI:** 10.1101/2021.02.04.429714

**Authors:** Mark A. Reimers, Kenneth S. Kendler

**Affiliations:** Department of Physiology and Institute for Quantitative Health Sciences and Engineering, Michigan State University, East Lansing, MI; Virginia Institute of Psychiatric and Behavioral Genetics, and Department of Psychiatry, Medical College of Virginia/Virginia Commonwealth University, Richmond, VA

## Abstract

We investigated the functional classes of genomic regions containing SNPS contributing most to the SNP-heritability of important psychiatric and neurological disorders and behavioral traits, as determined from recent genome-wide association studies. We employed linkage-disequilibrium score regression with several brain-specific genomic annotations not previously used. The classes of genomic annotations conferring substantial SNP-heritability for the psychiatric disorders and behavioral traits differed systematically from the classes associated with neurological disorders, and both differed from the classes enriched for height, a biometric trait used here as a control outgroup. The SNPs implicated in these psychiatric disorders and behavioral traits were highly enriched in CTCF binding sites, in conserved regions likely to be enhancers, and in brain-specific promoters, regulatory sites likely to affect dynamic responses. The SNPs relevant for neurological disorders were highly enriched in constitutive coding regions and splice regulatory sites. We suggest that our results provide a bridge between genetics and the well-known effects of life history and recent stressful experiences on risk of psychiatric illness.

## Introduction

Recent studies (e.g. (1)) have found that little of the SNP-heritability for psychiatric disorders lies in coding regions. These results provoke the question: what kinds of genomic elements are relevant to each psychiatric disorder – which we term the ‘functional genetic architecture’ of the disorder – and do the functional genetic architectures of psychiatric disorders differ systematically from those of complex neurological disorders or behavioral or anthropometric traits? By comparing the functional genetic architectures of psychiatric disorders to those of neurological disorders and behavioral traits, we sought to determine if the mechanisms of disorders differ systematically and how the resulting typology of illness relates to typology based on familial factors and/or SNP-based polygenic risk scores.

Twin and family studies have investigated the degree to which different psychiatric disorders share familial liability (2, 3). With the development of polygenic risk scores (PRS), evidence for substantial genetic correlations across various psychiatric disorders was replicated and extended (2, 3) while the correlations across psychiatric and neurological disorders were limited (4). These results are of interest outside the specialized area of psychiatric genetics because the familial/genetic relationships between psychiatric disorders are used as a primary method for clarifying nosologic boundaries between disorders (5).

However, a complementary approach to the genetic architecture of psychiatric and neurologic disorders examines the relative contributions of different functional classes of genomic elements, such as dynamic regulators, affecting response to experience, or constitutive regulators that may affect cell-type identity, coding regions etc. This is the approach taken here.

A separate important issue is whether the findings of psychiatric genetics can be integrated with the well-established findings of the life-history risk factors for mental illness (6, 7). Although psychiatric GWAS implicate many brain-related genes, especially synaptic genes, it remains unclear how the genetic risk factors may be related to the well-documented environmental risk factors for illness. A simple hypothesis is the that the genetic risk factors for psychiatric disorders lie predominantly in DNA that dynamically regulates genes in response to changing environmental circumstances or bodily signals, rather than in DNA that determines protein products or cell-type identity.

Several groups have attempted to partition the common variant (SNP) heritability of select psychiatric disorders among different functional categories. Schork et al (8) compared genetic contributions of different parts of coding genes and found that the untranslated regions accounted for more heritability than coding regions for schizophrenia; however, the authors noted that, because of the high linkage disequilibrium (LD) in the human genome, it is difficult to assign unambiguously a particular association signal to a particular SNP, and thereby to determine in which categories most heritability lies. This assignment is especially challenging for functional classes that are frequently juxtaposed on the genome, (e.g. transcription start sites (TSS) and promoters) so that SNPs in LD with a SNP in one functional class are often in high LD with a SNP in another class. Schork et al (8) attempted to resolve this ambiguity by adding all the annotations in LD with all SNPs of genome wide significance, weighted by the LD r^2^.

Finucane et al (9) addressed the issue of LD more systematically using partitioned linkage disequilibrium score regression (LDSR). This method exploits the wide distribution of risk SNPs with small effects and is based on the idea that SNPs in high LD with classes of SNPs most relevant to risk will have systematically elevated chi-square association scores. Their initial presentation used a large set of diverse annotations from different sources, including some regulatory types; they offered a preliminary assignment of SNP heritability among classes and found differences among traits. However, most of these annotations were not brain-specific, and significant improvements in the annotation of regulatory functions have been made since their use of generic ENCODE data. This is an opportune time to revisit the LDSR approach using more recent and brain-specific data.

The goals of this study are to characterize the functional genetic architecture of a range of psychiatric and neurological disorders and behavioral traits. We predicted that a preponderance of the heritability for psychiatric disorders and behavioral traits would be in regulatory sites, specifically enhancers, while most of the heritability for neurological disorders would be in protein coding regions. We further expected lncRNAs to contribute to psychiatric disorders because they were highly expressed in specific brain cell types and play critical roles during development (10).

## Methods

### 2.1 Sources of data

We annotated 9.5M SNPs in the human genome (HG19) as follows. We downloaded from the LDSR github site certain key generic (i.e. tissue-independent) annotations (e.g. coding regions) used in (9). We added selected several non-coding generic annotations from ENSEMBL, conservation data from UCSC and we included some brain-specific regulatory annotations based on chromatin data from RoadMap Epigenomics (11) and from PsychENCODE (12, 13). These annotations and their sources are summarized in Table 1.

**Table 1.** Genome annotations used in this study and their sources

| Annotation | Source | Reference | Comment | Proportion of SNPs |
| --- | --- | --- | --- | --- |
| Promoter UCSC | LDSR | Finucane |  | 0.0463 |
| TSS | LDSR | Finucane |  | 0.0178 |
| Protein coding | LDSR | Finucane |  | 0.0143 |
| 3' UTR | LDSR | Finucane |  | 0.0036 |
| 5' UTR | LDSR | Finucane |  | 0.0055 |
| Splice donor | Constructed |  | 70 nt from start of intron and conserved | 0.0024 |
| Splice acceptor | Constructed |  | 70 nt from end of intron and conserved | 0.0019 |
| Brain Promoter | RoadMap | RoadMap Epigenomics |  | 0.0031 |
| Mammal Conserved | UCSC |  | excluding other annotations | 0.0059 |
| Primate Conserved | UCSC |  | excluding other annotations | 0.0136 |
| CTCF binding | PsychENCODE |  |  | 0.0194 |
| lncRNA | ENSEMBL |  |  | 5.00E-04 |
| micro-RNA | ENSEMBL |  |  | 6.40E-05 |
| ribosomal RNA | ENSEMBL |  |  | 8.90E-06 |

We generated two new kinds of annotations. One often overlooked source of regulatory variability are splicing regulatory sites. These are commonly found on either side of the splice junction, but they are poorly known or annotated. We assigned SNPs provisionally to these categories if they were located on introns within 70bp of an annotated splice junction and conserved across mammals.

Since we expected much of the heritability of psychiatric disorders to be in regulatory regions such as enhancers, we gathered and used annotations of enhancers from several sources, based on chromatin assays. However, although annotated enhancers (based on H3K27ac or ATAC chromatin peaks) from these studies showed significant enrichment among SNPs implicated by psychiatric and behavioral GWAS, none explained more than 20% of SNP-heritability in the LDSR model. Some reasons for this are discussed below.

We adopted the following strategy to identify probable enhancers. Our annotation classes included all the known specific non-coding elements of the genome, many of which are highly conserved. We reasoned that most of the remaining non-coding regions highly conserved across mammals (PhastCons > 0.5) were likely to be enhancers, even though not all would be active in the brain. One well-known problem with using conserved regions to identify enhancers is that enhancers are typically not well conserved across different orders of animals; furthermore there has likely been recent rapid evolution of regulatory sites affecting the human brain. This problem was partially addressed by using primate conservation data from UCSC; only 20% of these primate-conserved regions overlapped other mammal-conserved regions, consistent with the rapid evolution of brain enhancers in the primate lineage.

### 2.2 Class-Specific Heritability Estimates

We used the LDSR procedure software provided by the Broad Institute (https://github.com/bulik/ldsc), and made the following modifications, both in line with their recommendations. First, two regions of very high linkage disequilibrium were excluded: the MHC region and the GPHN yin-yang region since both have strong associations with some psychiatric disorders and their leverage points would distort the regression. Second, the LDSR regression model tacitly assumes that all effect sizes within a category are comparable. However, the actual distribution of effect sizes observed in GWAS is very strongly right-skewed and outliers can substantially distort least squares fits, such as those used in LDSR. We therefore winsorised the summary P values at 10^−7^, corresponding to a chi-square of 22.

Besides the categories reported here we also used several other annotations of non-coding RNAs (microRNAs, and ribosomal RNAs). The proportions of SNPs with each of these annotations were less than 1 in 10,000, and the standard errors of heritability estimates for those classes from LDSR were almost all larger than the estimates and thus were omitted from the presentation.

Genome build made little difference to the results. Running LDSR for partitioned heritability on the same GWAS summaries using LD from either HG19 or HG38 had minimal impact on the heritability estimates. Since most of the GWAS results used here were reported initially in HG19, we used LDSR on this older build.

We obtained GWAS data from 18 brain-related phenotypes as listed in Table 2. We attempted to sample broadly from psychiatric disorders and behavioral traits (14-25), as well a selection of neurological disorders (26-30). We included well-studied biometric traits, height and BMI, as controls.

**Table 2.** Sources of GWAS data used in this study

| Trait | Acronym | Reference | Total N |
| --- | --- | --- | --- |
| Age-related Macular Degeneration | AMD | Fritsche et al. 2013 | 77255 |
| Alcohol Use Disorder | AUD | Walters et al. 2018 | 46568 |
| Alzheimer's disease | AD1 | Jansen et al. 2019 | 455258 |
| Alzheimer's disease | AD2 | Kunkle et al. 2019 | 63926 |
| Attention Deficit Hyperactivity Disorder | ADHD | Demontis et al.. 2019 | 55374 |
| Autism Spectrum Disorder | ASD | Grove et al. 2019 | 46350 |
| Bipolar Disorder | BPD | Stahl et al 2019 | 41653 |
| Body Mass Index | BMI | Yengo et al. 2018 | 681275 |
| Educational Attainment | EDU | Lee et al 2018 | 766345 |
| Epilepsy | EPI | ILAE, 2018 | 44889 |
| Extraversion | EXT | Van Den Berg et al 2015 | 63030 |
| Height | HGT | Yengo et al. 2018 | 693529 |
| Intelligence | IQ | Savage et al. 2018 | 269867 |
| Major Depressive Disorder | MDD | Wray et al. 2018 | 480359 |
| Neuroticism | NEU | Nagel et al.. 2018 | 380506 |
| Parkinson's disease | PAR | Nalls et al., 2019 | 482730 |
| Reaction Time | RT | Davies et al., 2018 | 282014 |
| Risky Behavior | RSK | Linner et al. 2019 | 466571 |
| Schizophrenia | SCZ | Pardinas et al 2018 | 105318 |
| Subjective Well-being | SWB | Okbay et al 2016 | 298420 |

The LDSR program was downloaded in March 2019 and run using recommended settings. The LDSR estimates are unbiased, thus the LDSR method yields some negative heritability estimates when the standard error of the estimates exceeds the (positive) true h^2^. The proportion of negative estimates of proportions of h^2^ was consistent with what would be expected if one third of the categories contributed much lower SNP-heritability than the standard errors of the estimates. These negative estimates occurred mostly for those traits, which themselves have low SNP-heritability (mostly behavioral traits).

LDSR estimates for some categories had standard errors within a factor of two of the estimates. In order to reduce the error, we used an empirical Bayes (eBayes) approach. We started by observing that for annotation classes with well estimated heritabilities, (i.e. small standard errors), the estimates followed an approximately exponential distribution across different phenotypes. Therefore, we modeled the distribution of h^2^ across phenotypes by an exponential for all annotation classes. We estimated the parameter for each class by maximum likelihood: we determined the exponential parameter that gave the highest probability for observing the full set of heritabilities estimated by LDSR across all phenotypes, taking into account the standard errors of these estimates (process documented in accompanying code). The posterior distribution of the estimate for each phenotype was then the exponential prior multiplied by the likelihood function, and the posterior estimates were computed as the expected value of the posterior distribution.

Empirical Bayes approaches introduce a bias in order to reduce unmodeled error. Since the aim of this paper is to document distinctions among phenotypes, and the bias of eBayes draws estimates for each phenotype toward the common mean of all phenotypes, the bias does not contribute to our results. We also tried a shrinkage strategy analogous to that used by the LASSO and found only very modest differences in results (not reported).

## Results

The partitioned heritability estimates for the most significant categories and the enrichments (ratio of proportion of SNP-heritability to proportion of SNPs) for selected categories are shown in Fig 1; the raw estimates from running the Broad LDSR program and their standard errors are presented in S1 Table. The classes contributing most to SNP-heritability were coding regions and transcription start sites (TSS; for most neurological disorders) and CTCF sites (psychiatric and behavioral phenotypes). The most enriched classes (contributing much more than their proportion) were these three classes and also brain-specific promoters (mostly for psychiatric and behavioral).

**Fig 1.**
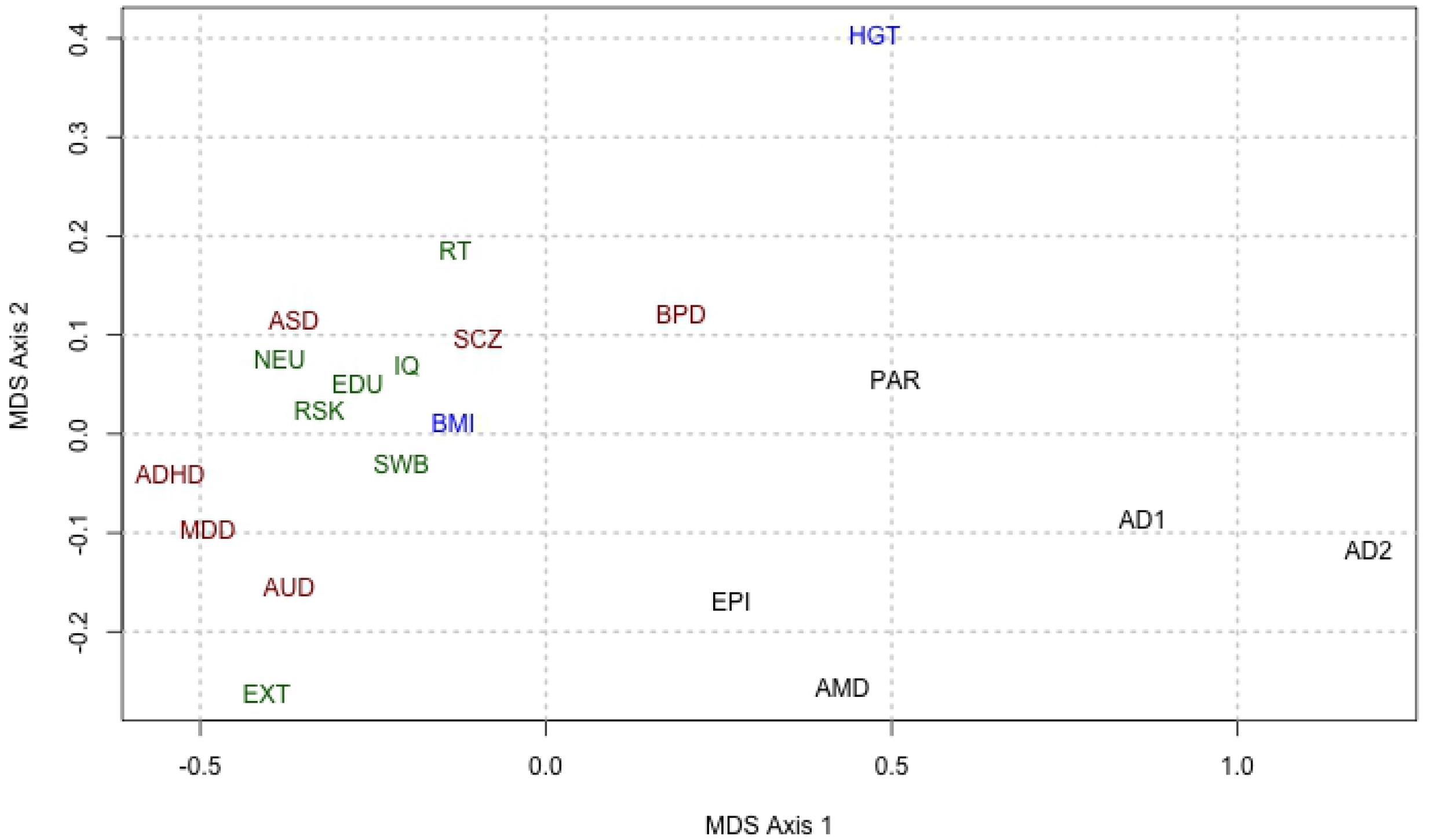
Heritability and enrichment estimates for 20 brain phenotypes. a) Empirical Bayes heritability estimates for the genomic classes studied here (in columns) for 20 traits and disorders (in rows). Color (legend at right) indicates estimated proportion of SNP-heritability. Estimates are (slightly) biased downward. b) Empirical Bayes enrichments of estimated SNP heritability attributed to various genomic classes by LDSR. Color indicates the enrichment (ratio of attributed heritability to proportion of SNPs) for each genomic category for each trait; key at right: blue: 0 (depletion); teal: little enrichment (1-2-fold); red: high (> 12-fold) enrichment.

The patterns of partitioned heritabilities seen in Fig 1 segregate with *a priori* classifications of the phenotypes, so we asked how the genetic architectures of the different traits relate to each other. We represented the relations among partitioned heritability patterns of phenotypes (Fig 2) using Kruskal’s isometric multi-dimensional scaling (implemented as isoMDS in R3.3) We defined distance between phenotypes by the sum over categories of the absolute differences in estimated heritability. The heritability distribution patterns of the core psychiatric traits cluster together with behavioral traits at center-left, while neurological disorders are spread through the lower right.

**Fig 2.**
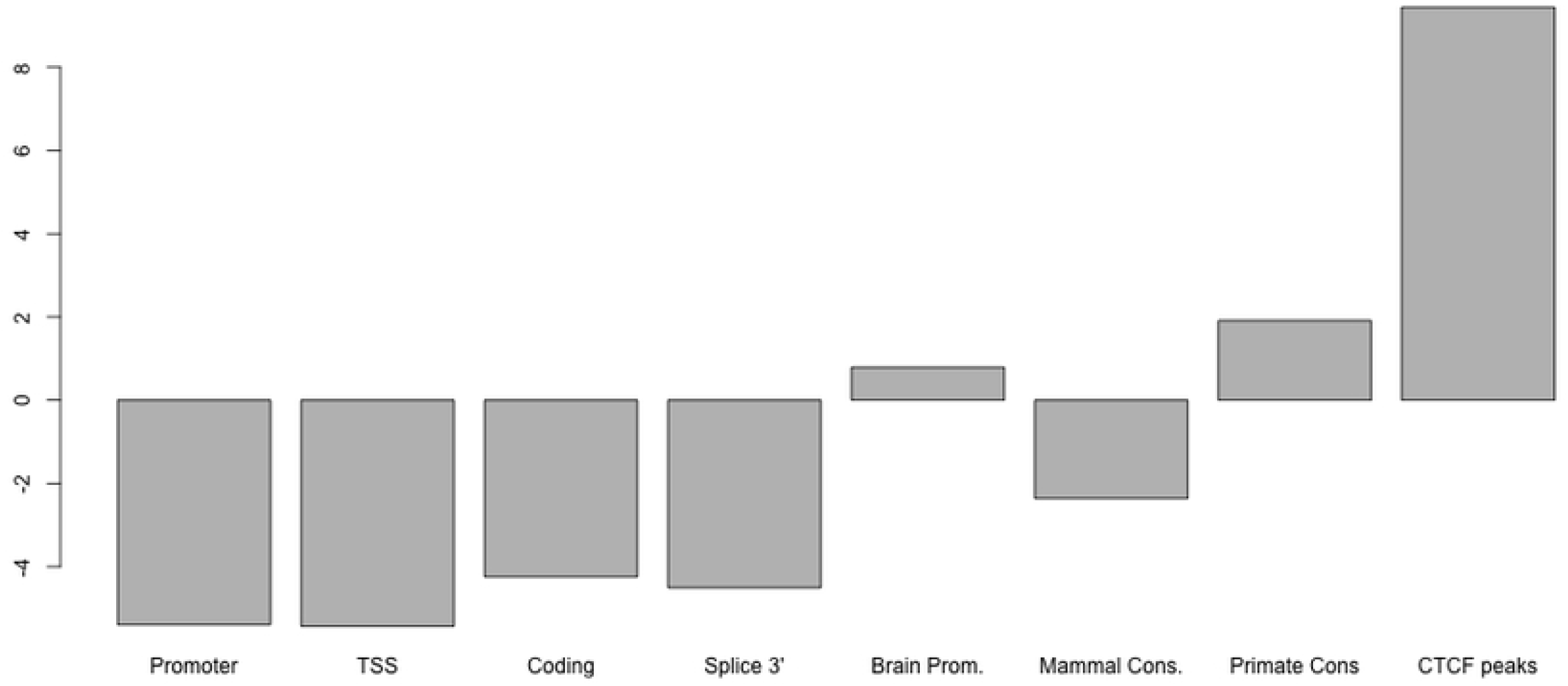
Multi-dimensional scaling 2-D plot showing similarities of functional genetic architecture among different traits. The horizontal axis corresponds roughly to higher loadings on constitutive (coding, promoter, splicing) annotations toward the right and higher regulatory related loadings toward the left. KEY: (for references see Table 2)AD1/2 Alzheimer’s disease (see Table 2); ADHD: Attention Deficit Hyperactivity Disorder; ASD: Autism Spectrum Disorder; AMD: Age-related macular degeneration; AUD: Alcohol use disorder; BMI: Body mass index; BPD: Bipolar disorder; EDU: Educational Attainment; EPI: Epilepsy; EXT: Extraversion; HGT: Height; IQ: Intelligence quotient; MDD: Major depressive disorder; NEU: Neuroticism; PAR: Parkinson’s disease; RSK: Risky Behavior; RT: Reaction Time; SCZ: Schizophrenia; SWB: Subjective well-being;

The clustered arrangement of traits in Fig 2 suggests that the partition of heritability among classes might be robust enough to distinguish whether an unknown disorder was neurological or psychiatric. To test this rigorously, we fit a linear discriminant to the heritability partition vectors and performed leave-one-out cross-validation. The predicted out-of-sample classes were the same as actual classes in all cases, confirming that patterns of enrichment can help distinguish between neurological and psychiatric disorders. Fig 3 shows the loadings of the discriminant function. The contribution of CTCF sites is the most discriminating measure, followed by contribution of coding regions (negative) and of primate-conserved regions. We were unable to find a robust linear discriminator based on genomic classes between behavioral traits and psychiatric disorders.

**Fig 3.**
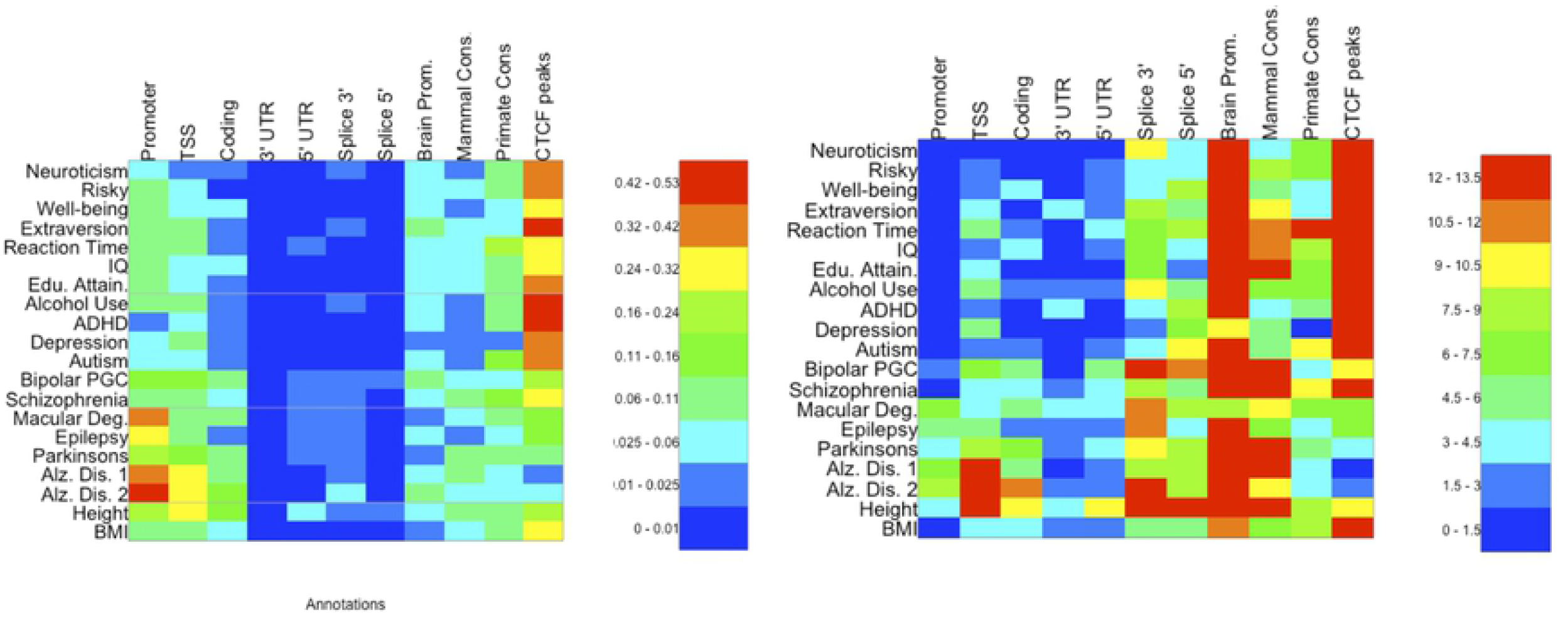
Functional genomic features that discriminate psychiatric disorders from neurological disorders. Bar plot showing weights of the linear discriminant function separating SNP functional class enrichment profiles typical of psychiatric disorders and behavioral traits (positive enrichments) from those profiles typical of neurological disorders (negative enrichments). Note heavy weighting on CTCF sites and putative primate enhancers for psychiatric disorders, but on coding regions for neurological disorders. Note that because the proportions of different SNP classes vary by almost three orders of magnitude, the discriminant weights displayed here were determined for enrichment ratios (heritability for a class divided by proportion of SNPs in that class) rather than heritabilities.

## Results Summary

We found that the majority of heritability for psychiatric disorders seems to be in putative regulatory sites: enhancers and CTCF sites. The sum of estimated SNP-heritabilities over all categories was similar for most traits: between 80% and 90%. These results suggest that the categories used here, although comprising less than 13% of the common SNPs in the genome, account for most of the SNP-heritability of these disorders or traits. Furthermore at least half the SNP-heritability for psychiatric and behavioral phenotypes seems to lie in less than 3% of the genome.

Notably we have found that brain-specific promoters and two relatively unstudied categories – CTCF binding sites, and putative inducible or cell-type specific enhancers – provide the majority of the SNP heritability for the major psychiatric disorders (schizophrenia, autism and bipolar disorder) as well as for behavioral traits, but not for neurological disorders.

We found that the three classes of non-coding RNAs tested (miRNAs, lncRNAs, rRNAs) appeared greatly enriched (medians 20-fold, 12-fold, and 20-fold, resp.) for contributions to SNP-heritability. Nevertheless, the overall contributions to heritability were small because the number of SNPs in such genes was small. Furthermore, the estimates of contribution were uncertain. Therefore, results from these categories are not presented. We have also identified a new category – putative splice regulatory sites – as relevant to neurological disorders.

## Discussion

We sought to determine whether we could distinguish the functional genetic architectures of psychiatric disorders, behavioral traits and neurological disorders. We predicted that variation in regulatory sites would play a greater role in the etiology of psychiatric disorders and likely behavioral traits than in neurologic disorders, while the reverse pattern would be observed for coding sequence variation. Our results partially confirmed these expectations.

Results for non-coding RNAs are not shown in Fig 1 because the standard errors of estimates for all three classes were comparable to, and usually bigger than, the estimates. Their presence or absence made little contribution to the relations among phenotypes or the appearance of Fig 2. We expected substantial heritability for psychiatric syndromes in long non-coding RNAs (lncRNAs) expressed during development. Indeed, we found that all classes of non-coding RNAs appeared enriched across all phenotypes, consistent with the emerging idea that non-coding RNAs play a role in human disease. Nevertheless, the estimates of SNP-heritability were all quite modest (under 1%), the LDSR standard errors were larger than the estimates in most cases, and differences between estimated enrichments across phenotypes or classes were not significant. However, all phenotypes with high estimates (at least ten-fold) for contribution of lncRNAs were behavioral or psychiatric; and for autism spectrum disorders the estimated proportion of heritability due to lncRNAs was greater than 1% and larger than two standard errors. Greater genetic resolution of GWAS may allow us to gain insight into role of these non-coding elements.

We were surprised to find such a strong representation of brain-specific CTCF sites in psychiatric disorders and behavioral traits, but only very modest enrichment (not shown) for the ENCODE CTCF sites used in (9).

Two of our new brain-specific categories – brain-specific promoters, determined from RoadMap Epigenomics data; and CTCF binding sites, determined from PsychENCODE data – contributed substantial heritability to psychiatric disorders and behavioral traits. The generic cross-tissue versions of these categories used by (9) did not contribute substantially to psychiatric disorders, although the generic promoters did contribute to neurological disorders (Note that the UCSC promoter annotations used by LDSR enclose more than ten times as many SNPs as the RoadMap brain promoters). Many genes have several promoters which may be active in different tissues. Use of different promoters will result in different 5’UTRs, which contain regulatory signals often related to trafficking the RNA to specific cell compartments, such as dendrites. The greater enrichment of brain-specific promoters and CTCF sites validates our rationale for using regulatory sites derived specifically from brain chromatin data, with one significant exception: enhancers.

The function of CTCF in gene regulation is poorly known and is now an active area of research. We do know that brain-specific CTCF binding sites are highly conserved across mammals, and hence must play an important role in the genome. Recent evidence suggests that CTCF and its induced chromatin looping are not required for basal cell functions. Rather, chromatin configuration is highly dynamic at the fine scale, and CTCF plays a major role in these reconfigurations, acting to stabilize DNA loops during enhancer-promoter contact on time scales of minutes (6). These findings suggest that CTCF binding sites are likely candidates for modulating dynamic responses to transient cell signals. In the brain, transient cell signals mediate learning (7). Some reports (31) indicate that CTCF plays a critical role in the brain’s most specific functions, such as learning and memory, CTCF would be strongly implicated in psychiatric disorders and behavior traits, but less so in neurologic disorders or biometric traits.

We identified brain promoters and expected to identify brain enhancers using data from published chromatin assays of human brain tissue (11). Brain promoters selected from the annotations produced by seemed useful, but, LDSR did not find that enhancer annotations from these data sets explained a large fraction of SNP heritability. We suggest two main reasons for this. First, currently available brain chromatin data is derived primarily from dissected tissue, aggregating across nuclei from all major cell types. Second, enhancers are not always active: many enhancers, especially those critical for learning are induced in only a small fraction of cells by specific signals and are ‘on’ for brief periods during which a burst of transcription is activated.

Enhancer annotations derived from the chromatin data currently available are thus likely to reflect predominantly constitutive enhancers in the most abundant cell types. Our success in finding enrichment signals in putative regulatory sites flagged by conservation, and our failure to find as much in chromatin data, suggests that inducible enhancers that are responsive to physiological signals and events or enhancers in minor cell types contribute to the genetics of psychiatric disorders more than other disorders. This interpretation is consistent with evidence that i) interneurons (32) and ii) physiological insults such as injury or infection or life experience stress (33) are implicated in psychiatric disorders more than in neurological disorders or biometric traits.

We were surprised to see that the functional genetic architecture of BMI seemed more similar to that of behavioral traits than to the standard biometric trait of height. However (34) found many SNPs relevant to BMI in or near genes expressed in the nervous system.

Does the relationship between psychiatric disorders assessed from our functional genomic categories seen in Fig 2 map onto those obtained from common SNP variants formed into polygene scores? A definitive answer is not yet possible, but two lines of suggestive evidence can be derived from the magnitude of SNP-based genetic correlations that bear some resemblance to the distance between the disorders in Fig 2. First, using SCZ as an anchor point, SNP-based genetic correlations are high between SCZ and BPD (positioned closely together Fig 2) and modest with MDD (which is further apart) (2). Second, using AUD as an anchor, SNP genetic correlations are high with MDD and modest with SCZ and BPD (35).

The results presented here complement recent studies showing genes implicated by GWAS for neurological disorders concentrate in specific brain cell types, while genes implicated by GWAS for psychiatric disorders and behavioral traits are broadly enriched in telencephalic neurons (36).

## Conclusion

In a novel use of LDSR, we have identified the genomic categories accounting for a majority of the SNP heritability for a number of major psychiatric disorders. We have also shown that the functional genetic architectures of many psychiatric disorders and behavioral traits are relatively similar to each other and less similar to the architectures of neurological diseases or to a control anthropometric trait like height. We have shown that distinctive genomic categories relevant to psychiatric disorders and behavioral traits are those related to dynamic gene regulation on short time scales. Our results hold promise for bridging genetics and well-established environmental and life-history risk factors for psychiatric disorders.

## Acknowledgments

This project was supported by a Genetics and Human Agency Award to KSK. This work is solely the responsibility of the authors. and does not necessarily represent the official view of the funders (John Templeton Foundation). We thank Amanda Charbonneau and Jorden Schossau for technical work with LDSR, and Bradley Verhulst for consultations about the GWAS used here.

## Supporting information

**S1 Table. LDSR Results**. These are the results of running the LDSR program with the options noted in the text.

